# DNAS-Bench: Deterministic Nucleic Acid Screener Benchmarking

**DOI:** 10.64898/2026.07.06.736904

**Authors:** Henry C. Wong, Tadayoshi Kohno, Jeff Nivala

## Abstract

The rapid growth of biotechnology manufacturing for synthetic DNA and proteins has raised concerns that adversaries could exploit commercial synthesis pipelines to create biological weapons. Without effective safeguards, an attacker could seek regulated genetic sequences from synthesis providers; while synthetic DNA is not itself a pathogen or toxin, access to such sequences can lower barriers to downstream misuse, motivating robust order-time screening. To mitigate this risk, Biosecurity Screening Software (BSS) systems have been developed to flag potentially malicious synthesis orders. Here, we propose one of the first deterministic benchmarks for evaluating the robustness of Biosecurity Screening Software. Our framework enables systematic testing of BSS behaviors and potential on specific nucleic-acid sequences and on targeted regions of malicious genomes. Our framework allows for insights into what is being flagged as malicious in BSSs, leading to potential discussions if specific BSS is fit for a specific manufacturing pipeline. We additionally introduce a dataset of manipulated genomes derived from the HHS and USDA Select Agents and Toxins List. When evaluated on this dataset, SeqScreen flags 42% of the sequences as malicious, while Commec flags 10.2%. Across a range of manipulation strategies, we find that simple manipulations, such as padding sequences by adding a repeated nucleotides at 1.5 times the original length, perform nearly as well as more targeted methods, such as embedding malicious sequences within benign genomic context. Padding-based methods trail embedding-based methods by only 0.75 percentage points in average detection rate. Consistent with prior reports from BSS developers and studies, we observe a sharp drop in detection rate when input sequence length falls below a critical threshold, typically between 50 and 100 base pairs (bp). Under our threat model, this implies that an adversary can bypass most existing safeguards by splitting a target genome into fragments shorter than 50 bp. Fragment-level analysis further reveals that some toxin regions evade detection entirely by SeqScreen, while other malicious genomes remain detectable even when fragmented into 30–50 base-pair segments. We open-source this benchmark to support reproducible evaluation of BSS robustness and to inform the development of next-generation biosecurity screening tools (https://github.com/HenryCWong/DNAS-Bench). For ethical concerns we only open-source the framework while the data is available upon request.

## 1. Introduction

One of the growing concerns in cybersecurity and biology is the misuse of rapidly progressing biotechnology, introducing new possibilities for bioterrorism and the creation of new bioweapons. Advancements in synthetic biology allow customers to order synthetic nucleic acids online and receive their orders within a week. Customers can then process the nucleic acids to make proteins, engineer organisms, develop drugs, and much more. An adversary can potentially pose as a customer and order malicious nucleic acids, such as sequences from a virus like influenza, and process the nucleic acids into a bioweapon.

These advanced capabilities have raised concerns over potential misuse [1] leading to conversations of regulation by the United States government [2] [3] [4] [5]. In 2024, the U.S Office of Science and Technology (OSTP) directed all federal agencies funding life sciences research to screen all synthetic DNA orders to a certain standard [6]. Some synthetic DNA companies already have software systems to screen orders called Biosecurity Screening Software (BSS). In the past few years, both commercial (Ultraseq [7], Fast-NA [8]) and open-source (Common Mechanism [9], SeqScreen [10]) tools have emerged. However, previous work has shown consumers can still order malicious sequences [11] and there have been recent studies on the use of generative AI to bypass BSSs [12]. The International Gene Synthesis Consortium, an industry-led organization established to promote biosecurity screening, also has its own guidelines for screening [13]. Individual DNA synthesis companies, like Twist Bioscience, have also published concerns about and work on screening [14]. To help improve the state of BSSs, we introduce DNAS-Bench. DNAS-Bench evaluates the screening efficacy of BSSs by pitting BSSs against adversarial generated datasets and examining the results through a suite of analytics tools. Our work is the result of cross-disciplinary collaboration between researchers in computer security and researchers in biotechnology. We introduce a deterministic benchmarking framework for BSSs using crafted adversarial datasets based on manipulations we believe a motivated attacker might perform.

In our threat model, we prioritize transformation methods without Artificial Intelligence involved due to the lack of deterministic studies and current BSSs struggling to handle deterministic manipulation methods. We also introduce more complex deterministic methods made to either mislead or circumvent recent patches to BSSs.

To demonstrate our benchmark, we test our dataset with two open-source BSSs and one taxonomic labeling tool. We take known sequences with Sequence-of-Concerns (SOCs) and split them into different lengths (ranging from 50 nucleotides to 300 nucleotides). Each following manipulation is a variation of multiple fragments of a sequence. When splitting a sequence into multiple fragments without any manipulations on the fragments, SeqScreen flags 75% of the fragments when the each fragment length is 300 base pairs while Commec flags 34.91%. This detection rate drops to 52.86% for SeqScreen and 20.12% for Commec when a random sequence 1.5 times the size of the fragment is appended to the end of the fragment.

Our results highlight the need for more security evaluations and adversarial testing on BSSs. We hope the community responds by putting greater effort into developing more robust open-source BSSs and constructing additional benchmarks similar to ours. Additionally, by developing our benchmarking framework and making it available to BSS designers and the security research community (we opensource our framework and processes and provide malicious data on request, see Appendix A). On our ethical considerations and plans for sharing our software, we hope to empower BSS designers with the ability to better evaluate the efficacy of our systems. Additionally, making our framework available to DNA synthesis companies, we make it possible for them to better assess the risks by using their selected BSS software.

### Summary

Our four key contributions include:

- We create a new benchmarking framework for testing Biosecurity Screening Software using a range of deterministic modifications.
- We evaluate two BSSs and one genomic classification model against a widely used select toxins and agent’s dataset.
- We derive a threat model for synthetic nucleic acid manufacturing.
- We make the benchmark available via open source for BSS developers for evaluation and future improvements to BSSs.

### Paper Structure

We begin with background (Section 2) in current work in cybersecurity in synthetic biology. In Section 3, we outline our threat model, misuse of a synthetic nucleic acid or protein manufacturing process, and the type of attack we analyze in this work. Section 4 provides an overview of the exact types of manipulation we perform on nucleic acid/protein orders. Section 5 shows our experimental setup of which BSSs we review, how we set up each BSS, and the data we choose to manipulate to test each BSS. Section 6 reveals the results of our experimentation which we discuss in Section 7.

## 2. Background

### 2.1. Synthetic Nucleic Acids and Proteins

Synthetic nucleic acids are artificially manufactured DNA or RNA sequences that are chemically synthesized rather than isolated from natural biological sources [15], [16]. Unlike traditional molecular cloning, synthetic nucleic acid production does not require a template, allowing researchers to design and construct virtually any desired sequence [16]. These synthetic molecules can serve a wide range of applications across life sciences such as primers and probes for polymerase chain reaction (PCR) assays and molecular diagnostics, as components of gene therapies and DNA vaccines, as tools for genome editing (such as CRISPR guide RNAs), and as building blocks for constructing synthetic genes, genetic circuits, and even entire genomes in synthetic biology research [15], [17].

The commercial genomic synthesis industry has transformed modern molecular biology by enabling researchers to obtain custom DNA sequences without the need for traditional cloning methods. Modern gene synthesis relies on the chemical synthesis of short oligonucleotides (typically under 200 base pairs) using phosphoramidite chemistry, followed by enzymatic assembly into larger constructs using methods such as Gibson Assembly [18] or polymerase cycling assembly [16]. These advances have enabled the construction of not only individual genes but also genetic pathways and even entire synthetic genomes [15], [17]. The development of commercial DNA synthesis services means that researchers can now simply upload a desired sequence to a provider’s website and receive synthesized DNA within days, often at costs below $0.10 per base pair [16]. This accessibility has accelerated research in synthetic biology, therapeutics, and biotechnology, but has also raised biosecurity concerns [19]. In response, the International Gene Synthesis Consortium (IGSC) was established in 2009, and member companies voluntarily screen double-stranded DNA synthesis orders over 200 base pairs against databases of regulated pathogens and sequences of concern [19].

### 2.2. Security Concerns in Synthetic Biology

Synthetic biology and biotech manufacturing have expanded considerably over the past few decades, with the market currently estimated at $16.35 billion and projected to reach $80 billion by 2034. However, the speed of this growing market has left many concerned about potential malicious actors and misuse. The United States Department of Health and Human Services (HHS) established recommendations for validating customer legitimacy and conduct sequence screening to flag orders containing Sequences of Concerns (SoCs) in 2010. And in 2024 the White House directed the Office of Science and Technology Policy (OSTP) to collaborate with several federal agencies to develop the “Framework for Nucleic Acid Synthesis Screening” [2], which requires recipients of federal funds to purchase synthetic nucleic acids from providers that follow the OSTP Framework. The framework requires providers and manufacturers to screen purchase orders for synthetic nucleic acids and report illegitimate purchases involving SoCs.

The National Security Commission on Emerging Biotechnology lists in their “Charting the Future of Biotechnology” report in Chapter 4.4 [3] the need to protect against the harms of biotechnology, calling for US congress to direct the executive branch to advance security biotechnology research and innovation, and specifically mentioning synthesis screening as a major concern.

### 2.3. Safeguards in Synthetic Biology

There has been prior work in outlining benchmarks for BSSs [6], which proposes a prototype dataset for evaluating DNA synthesis screening. However, that work does not apply adversarial manipulations to sequences (i.e., modifications an attacker might use to evade screening). The National Institute of Standards and Technology (NIST) has also released a dataset for assessing nucleic acid screening [20]. But, the work similarly lacks manipulated sequences and is relatively small (around 200kB). [21] performs an analysis of both commercial and open-source BSSs and finds that all tools had a baseline performance of 95%. However, the study does not consider manipulated data.

There has been previous work on using artificial intelligence to create novel formations of malicious nucleic acids and proteins [12]. This work describes an “AI red teaming” effort using open-source protein sequence generative models to create synthetic homologs of potentially malicious proteins. Working directly with BSS developers, the authors helped produce patches to commercial BSSs to improve the detection rate of AI generated malicious sequences. While patched, BSSs showed improved resilience to this manipulation compared to unpatched versions, detection remained imperfect, as both AI-based reformulation and deterministic manipulation reduce the length of contiguous sequence regions mappable to known proteins of concern.

In general there is concern of AI aided malicious use of synthetic DNA. In [22], concerns of dual-use risks in protein design and bioengineering AI tools are raised. AI companies like Anthropic and OpenAI have openly addressed attempts at filtering CBRN (Chemical, Biological, Radiological, and Nuclear) information [23] [24] and establishing red-teaming exercises, however few of these safeguards established are straightforward.

Deterministic manipulation of sequences has previously been explored [25] which describes most BSSs’ inability to correctly classify manipulated sequences by creating a dataset where non-coding DNA segments are inserted between toxin fragments. Similar to our methods, the authors also split sequences into fragments [25]. And ultimately finds that commercial tools can detect most non-manipulated toxins and have difficulty detecting splicing-based manipulations. Detection rate drops significantly once fragments are smaller than 50 base pairs. The work claims to have implemented a new algorithm in SeqScreen that has achieved 100% detection of manipulated sequences.

There are a handful of screeners like UltraSEQ [7], FAST-NA [8], Aclid, and NTI, all of which are proprietary and require payment. There are a select number of screeners that are open source like SeqScreen [10], Common Mechanism [9], and ToxinPred [26]. SecureDNA [27] offers a free, automated system that claims to block hazardous DNA synthesis orders as a part of a bigger service that includes cryptographic techniques to preserve customer privacy. SecureDNA also uses what they call a “Random Adversarial Threshold” that is designed for evasion resistance. Although SecureDNA’s source code is open source, the database is kept private. In this study, we focus on opensource screening. Open-source environments allow for better collaboration between researchers and developers and allow for more rigorous testing by the public [28].

## 3. Threat Model

Figure 1 illustrates our threat model. An adversary attempts to order a malicious sequence for purposes that could cause societal harm (e.g bioterrorism). At step **(1)**, the adversary finds a malicious sequence they want to use (e.g., influenza) and in step **(2)**, manipulates the sequence. In this threat model’s scenario, the adversary splits the sequence into multiple different fragments and in step **(3)** uses multiple thin clients and accounts to imitate the action of multiple users submitting different small fragments to synthetic nucleic acid manufacture, making detection through order patterns more difficult. In step **(4)**, before the synthetic nucleic acid is manufactured, the sequences must go through a BSS to ensure orders are not malicious. At step **(5)**, once the orders are confirmed non-malicious, the nucleic acids are manufactured. In step **(6)**, they are then sent back to the customer, in this case the adversary. Step **(7)** is where the adversary can assemble or manipulate all the fragments in a lab. Our study looks at how an attacker can manipulate and split their order at steps (2) and (3), then have those manipulated orders bypass the BSS at step (4) without issue.

**Figure 1.**
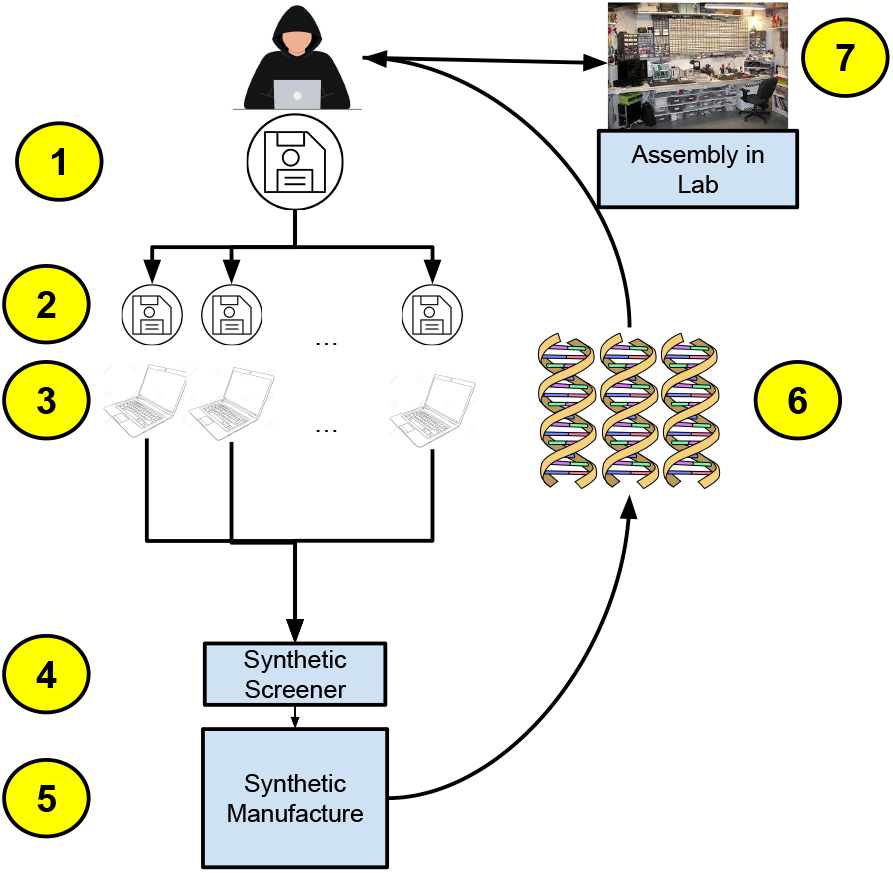
Threat Model

### 3.1. Attacker Capabilities

We restrict our attention to attacks without access to BSS code or infrastructure. Thus, we study an attacker with the following restrictions: no access to BSS code, no access to manufacturer’s computing infrastructure, and no physical access to the provider’s manufacturing process.

While sequence information is widely accessible and fragmenting text strings is straightforward, converting ordered fragments into a harmful biological outcome typically requires substantial laboratory capability (assembly validation, cloning/expression systems, appropriate host cells, troubleshooting, and verification) and may trigger regulatory and containment requirements depending on the agent/toxin. We therefore separate (i) feasibility of submitting sequences that evade automated screening from (ii) feasibility of downstream biological reconstruction, and we focus empirically on the former.

## 4. Benchmarking

The primary purpose of this paper is to design a challenge that not only tracks the performance of a BSS but also better defines where a BSS’s weaknesses are. In this section, we give an overview of our deterministic techniques. Each manipulation method we present does not require an attacker to have expertise in molecular biology knowledge. Rather, the attacker simply needs to have an adversarial mindset. Specific expertise is required only after delivery, when the attacker must assemble the fragments.

### 4.1. Manipulation Methods

Many regulated agents have genomes that are far larger than typical single-construct gene-synthesis products, and providers commonly deliver DNA as shorter fragments, oligo pools, or cloned segments rather than as complete genomes. Larger targets may be obtained only via multifragment synthesis and assembly, either by the customer or by the provider under additional controls. For benchmarking, we focus on fragmentation/manipulation patterns that stress screeners, while acknowledging that real-world feasibility depends on product type, provider policies, verification, and downstream assembly capability. We assume the attacker has access to standard assembly equipment available commercially. The splitting of a sequence can also service as a form of manipulation. We visualize some of the methods in Figure 2. We start thinking about these manipulations out of what is easiest for an adversary and then increase complexity from there. We explain details and provide pseudocode in Appendix B. Scripts used to split sequences are provided in the open-source Github repository.

**Figure 2.**
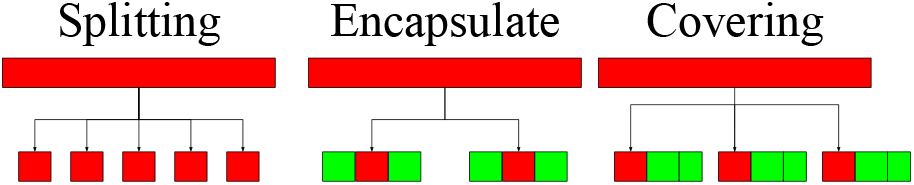
Visualization of manipulation methods to manipulate malicious agents

#### 4.1.1. Splitting

We split the malicious sequence into smaller fragments to give the screener fewer patterns to recognize, allowing the adversary to use assembly methods to piece together the sequence after receiving the physical nucleic acids. These fragments do not overlap each other and each numbered fragment is sequential in relation to the entire malicious sequence we download from NCBI. Other studies [25] on manipulating sequences include additional sequence elements a consumer may need for downstream activity like gene promoters, terminators, assembly adapters, and more. In this study, we do not add sequences such as these to study the direct effects of each manipulation.

#### 4.1.2. Encapsulation

We encapsulate each fragment with short flanking sequences of 4 base pairs as a simplified proxy for common ordering formats where fragments include additional bases beyond the core target (e.g., adapters, primers, or assembly-enabling context). This abstraction is not intended to faithfully model any single cloning method, but rather to test whether adding short non-target context around a fragment measurably changes screener behavior. This manipulation also represents a possible non-malicious action where the actor only intends to add assembly ends and is unaware that what they are ordering may be malicious. This can also be interpreted as a low-effort actor, only adding the minimum number of modifications needed. In this manipulation, we can also see how BSSs perform when the start and end of an order may not be malicious, rather than in Section 4.1.4 where the tail-end is only appended to.

#### 4.1.3. Flipping

We introduce controlled nucleotide perturbations as a benchmark stressor for sequence-matching approaches. Biologically, arbitrary mutations often destroy or attenuate function and restoring a functional agent after substantial sequence perturbation typically requires significant domain knowledge and additional laboratory steps. Accordingly, we treat this manipulation primarily as an algorithmic evasion analogue rather than a claim of low-effort real-world construction.

#### 4.1.4. Covering

We search for a benign sequence similar to the malicious sequence such that we can append or encapsulate the benign sequence to the malicious sequence in an attempt to force the aligner in the synthetic screener to align better to the benign sequence. The adversary can then remove the benign sequence after receiving the physical nucleic acid. This typically requires some mechanism like restriction enzyme digestion or PCR with primers designed to bind only within the target region. For our study, we do not include restriction enzyme sites as the intent of our dataset is to benchmark BSSs rather than create fully functional malicious sequences. Most DNA sequencing screeners use alignment software to attempt to pair the input sequence to an established reference sequence. If the size or importance of the benign sequence is significant enough, the screener could assume the malicious portion of the sequence is not as relevant and ignore the malicious sequence. In our study we use covering in three different forms. “Covers” is where we append a GFP gene sequence to the tail end of each fragment that is 1.5 times the length of the fragment. If the GFP gene is not long enough to cover 1.5 times the length of the fragment, we repeat the GFP gene until we reach the correct length. “Covers-a” is where we append a sequence composed only of adenine to the tail end of the fragment that is 1.5 times the length of the fragment. “Covers-random” is where we append a random genomic sequence that is 1.5 times the length of the fragment to the tail end of the fragment.

### 4.2. Pipeline

Our pipeline consists of three crucial steps that allow users to identify strong and weak points in BSS software through a suite of analytics. The first step, manipulating sequences, is outlined in Section 4.1. The second step runs a range of BSSs against all manipulations. In the third step the pipeline then generates analytics that allows users to view all agents that have been selected across different fragment lengths. Then to view a more in-depth analysis, the user can view how each fragment in a specific agent was viewed by two different BSSs along with annotations to show where each gene in the agent’s genome is relative to the results.

## 5. Benchmark Experimentation Setup

For this study, we reviewed eleven different agents from the HHS and USDA Select Agents and Toxins list [29] from the NCBI database that is publicly available [30]. To measure performance of our dataset, we compute the average classification detection rate (the rate at which the screener flags each fragment) among multiple splits for each select agent and task. We evaluate SeqScreen [10], Commec [9], and Kraken [31]. SeqScreen and Commec are both BSSs while Kraken is a taxonomic classifier. We benchmark Kraken for experimental purposes (see Section 6.2). For each model, we use the full database. For example, Common Mechanism only requires a 2 GB database; however, the full database including NCBI taxonomy and DIAMOND-formatted [32] databases requires almost 1 TB of data. For Common Mechanism we used the full 1 TB database. This is to ensure that each model is running at maximum potential to test our dataset. We also set up each screener with the default settings outlined in their published documentation.

We chose exclusively open-source screeners due to the importance of rigorous public testing for potential future work. Access to proprietary tools is difficult, expensive, and less transparent. We note that only SeqScreen and Commec are dedicated classifiers for malicious sequences while Kraken is a taxonomic identifier (see 6.2). We run all of our experiments on the latest version of all software as of November 2025 (SeqScreen version 4.5 and Commec version 1.0.1). Since the publication of our framework, Commec has released 4 new version to version 1.0.5 that includes many changes intended to increase accuracy and user experience.

We split each malicious sequence into consistent lengths from 300 to 10. For example, we run SeqScreen 291 times for a genome split into equal parts at fixed lengths between 300 to 10. We run SeqScreen where the length of each fragment of the genome is 300 base pairs (not including the assembly heads and tails), then 299 base pairs, and so on. We recognize that the very short fragment lengths considered here (e.g., 50 bp) do not reflect the most convenient or economical real-world synthesis workflow. We therefore frame these experiments as a conservative stress test rather than as a claim about the most likely adversarial implementation. Importantly, such short fragments are still technically assemblable using overlap-based gene assembly methods, although doing so becomes substantially more cumbersome and resource-intensive as fragment length decreases.

For splitting, each malicious sequence is split into fragments of a fixed length and put into a singular file. The other manipulation methods are similar but with their respective additions to each sequence. For example, in covering, a subsequence of a GFP gene is added until the GFP gene is 1.5x the length of the fragment. We then run each tool on every file. We then run each tool and file combination twice to ensure reproducibility.

### 5.1. Benchmark Dataset

We choose to use the HHS (US Department of Health and Humans Services) and the USDA (United States Department of Agriculture) Select Agents and Toxin list as a foundation for what malicious sequences to look for.

Out of the Agents and Toxins list we choose eight different toxins and agents to constitute our benchmark dataset. Given our ethics analysis, our intent is to provide this list of agents to researchers and others seeking to improve their DNA screening pipelines unless strong ethical arguments are presented to the contrary. Sequence 2 is separated into three different plasmids: Sequence 2.1, Sequence 2.2, and Sequence 2.3. Only Sequence 2.1 contains toxin genes, but Sequence 2.2 and Sequence 2.3 are still crucial for function. Sequence 8.1 and Sequence 8.2 are the same agent but were sequenced at over 10 years apart. We include these to evaluate whether the duration a sequence has been publicly available affects BSS performance.

### 5.2. Visualizations

We use visuals to support our analysis of BSSs’ classification rate through multiple adversarial-based manipulations. We first plot the detection rate of the BSS across each dataset for each type of manipulation (see Figure 3 and 5). Then, when a dataset is of interest, we look further into data by graphing every fragment across the genome of the dataset (see Figure 4). Figure 4 also allows for the option to add annotations so we can view exactly which genes are being classified. We can also compare BSSs with each other like in Figure 6 and create a comparison chart to compare multiple manipulation methods across a single genome.

**Figure 3.**
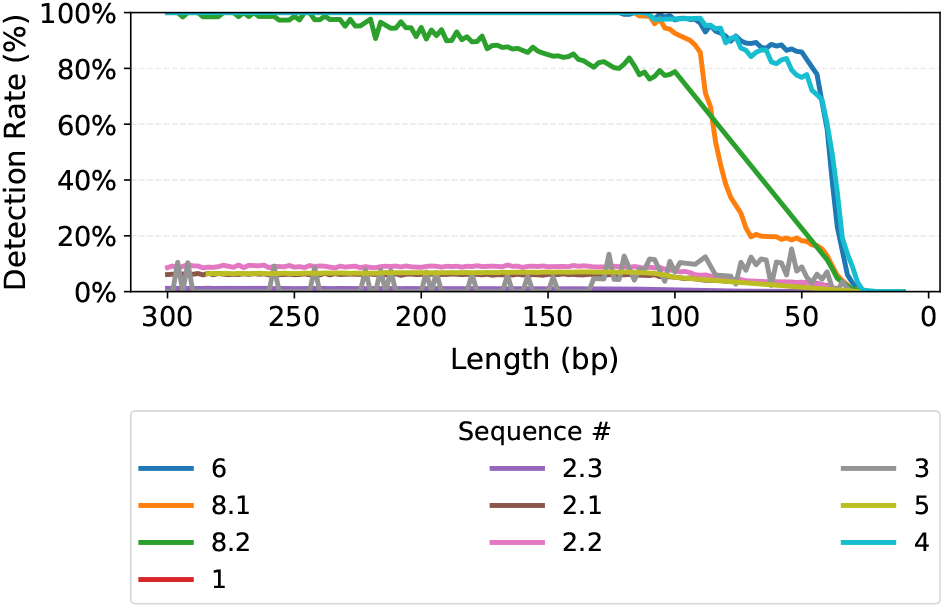
Detection rate at each encapsulation length for each agent in SeqScreen

**Figure 4.**
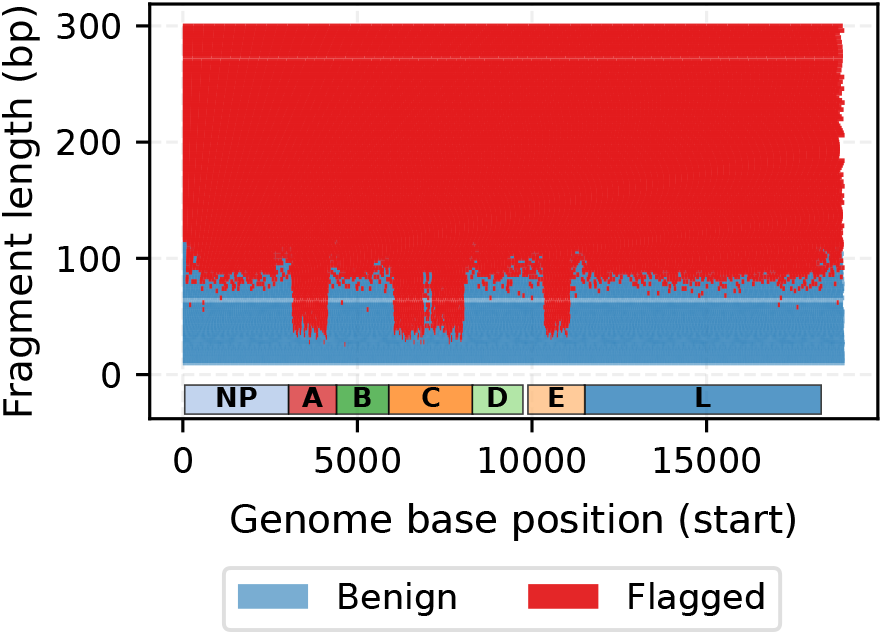
An in-depth overview of how SeqScreen performs against Sequence 8.1 across the entire genome where the genome is split at different lengths. The diagram along the x-axis shows positions of the different protein-coding regions within the genome.

**Figure 5.**
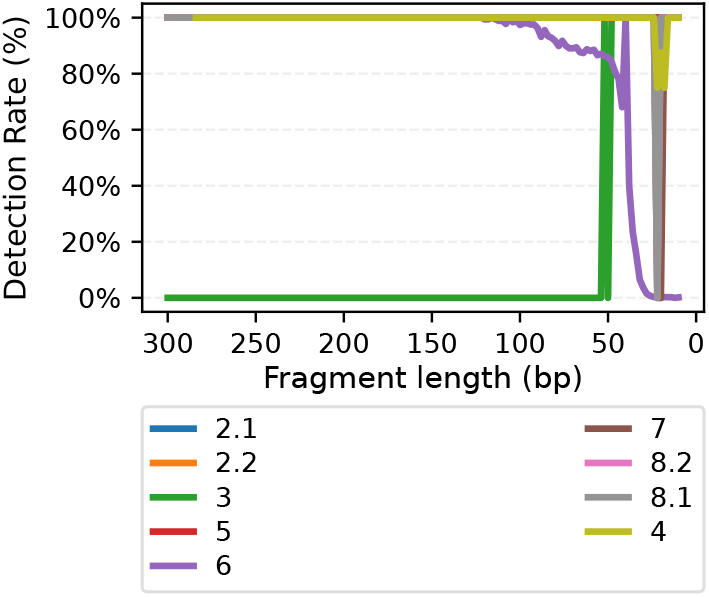
Detection rate of each tool on unmanipulated split-only datasets across fragment lengths for all sequences in SeqScreen.

**Figure 6.**
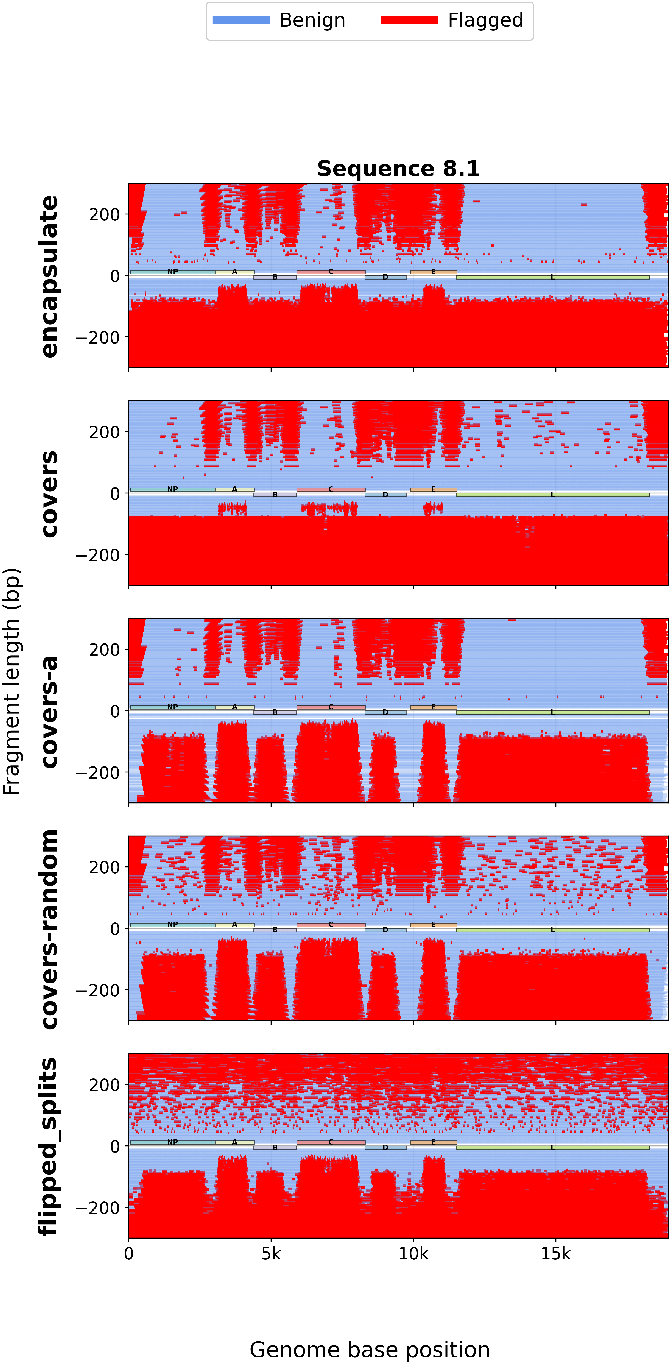
Comparison plots showing how SeqScreen and Commec perform in Sequence 8.1 with SeqScreen on the bottom half of each graph and Commec on the top half.

## 6. Results

In this section, we discuss the results of applying our benchmark manipulations and dataset to BSSs. In doing so, we demonstrate that our benchmark challenges today’s BSSs and offers a basis for future BSS benchmarking and evaluation.

In more detail, we evaluate two major open-source BSSs and Kraken, an informatics tool that only has the task of classifying sequences rather than determining if the sequences are toxic. Tables 2 and 3 give a summary of the performance between the two BSSs. We use the term “score” in Table 2. Since these numbers are computed over multiple manipulations and sequences from the benchmark dataset, we emphasize that the numbers are meant for comparison rather than a reflection of the individual BSSs’ performance in any specific use case. For our benchmark, SeqScreen flags more fragments than Commec by an average of 32 points and in every manipulation method we test.

**TABLE 1.** De-identified list of sequences used in experiments. Agent or toxin names are replaced with numeric IDs ; descriptions are intentionally high -level to avoid uniquely identifying specific isolates of constructs .

| Toxin or Agent (ID) | Description | Approximate Size |
| --- | --- | --- |
| 1 | Protein-coding toxin gene sequence (synthetic construct) | 753 bp |
| 2.1 | Bacterial virulence plasmid (large, DNA plasmid) | 181.9 kb |
| 2.2 | Bacterial virulence plasmid (large, DNA plasmid) | 94.27 kb |
| 2.3 | Bacterial chromosomal locus (DNA fragment) | 5.2 Mb |
| 3 | Bacterial pathogen genome segment (DNA) | 282.5 kb |
| 4 | Viral genome (segmented, RNA virus) | 13.6 kb |
| 5 | Bacterial pathogen whole genome (DNA) | 2.1 Mb |
| 6 | Viral genome segment (RNA virus) | 19.1 kb |
| 7 | Viral genome (RNA virus; alphavirus) | 11.7kb |
| 8.1 | Viral genome (RNA virus; historical isolate) | 19 kb |
| 8.2 | Viral genome (RNA virus; recent outbreak isolate) | 19 kb |

**TABLE 2.** Scores among all tested data of all Biosecurity Screening Software and Kraken in this study .

| BSS | mean | median | auc | mean at length=300 |
| --- | --- | --- | --- | --- |
| SeqScreen | 41.99 | 47.06 | 42.06 | 49.57 |
| Commec | 10.19 | 11.94 | 10.16 | 28.72 |
| Kraken | 40.08 | 45.95 | 40.18 | 49.77 |

**TABLE 3.**
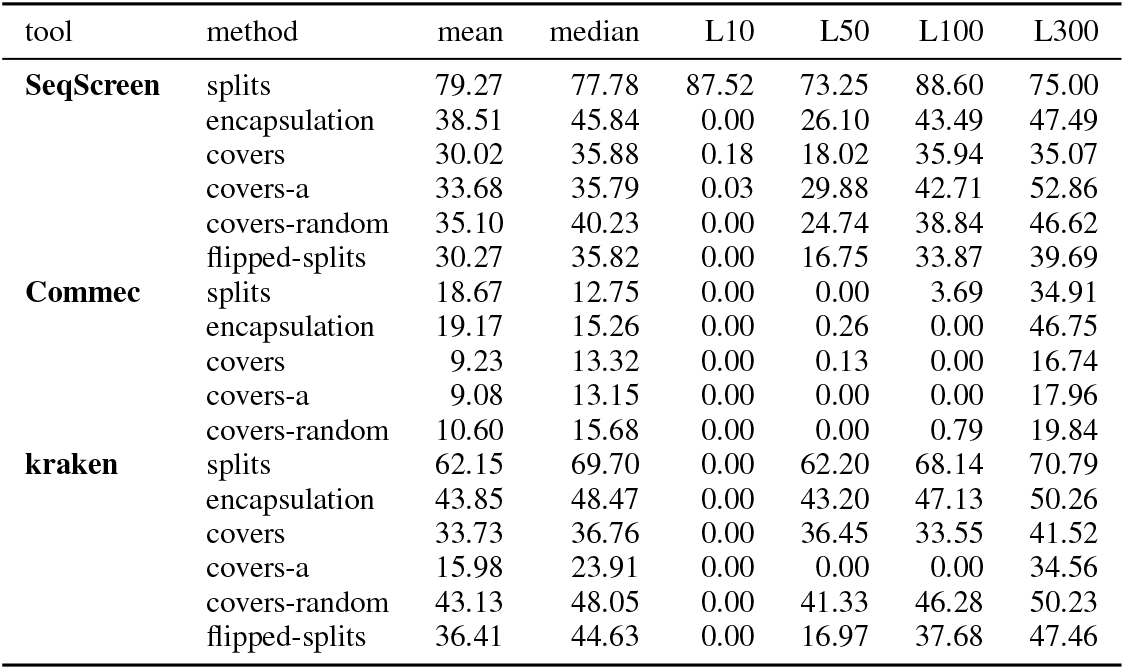
Average detection rate for each manipulation method for each BSS, with rates at selected fragment lengths. L300 indicates mean detection rate at all data when fragment length is 300 base pairs .

Figure 5 shows the results of SeqScreen when each dataset is simply split into multiple fragments with no manipulation to each fragment. SeqScreen can perform well on most datasets until the lengths of each fragment are less than 50 base pairs, with the exception of Sequence 3, whose detection rate increases as the fragment lengths get shorter. Interestingly, SeqScreen’s detection rate on a significant number of datasets dips at around 20 base pairs and comes back up soon after. However, we find that the classifications when SeqScreen starts flagging again, often classifies as the incorrect genome.

In Figure 3, SeqScreen attempts to classify malicious genomes that have been encapsulated, and we can visualize the performance drop suddenly when the length of the sequence reaches a specific point. There is a clear separation of datasets where SeqScreen can classify and cannot. For each dataset, each has a slightly different range where performance sharply drops, typically between the lengths of 50 base pairs and 10 base pairs of each fragment. Some agents, such as the Sequence 6, drop in detection rate at a length of 30. While there are groups of agents that do not perform well in general, such as Sequence 5 and Sequence 2.1.

Tables 5 and 6 show the detection rate of each respective tool’s rate at decrementing lengths of 50 from 300 to 10 base pairs for encapsulation data. These tables demonstrate exactly how performance decreases for each dataset as fragment lengths get smaller. Commec is the only BSS that can classify Sequence 1 despite not flagging other datasets. The tables also better demonstrate a consistent drop off in performance of all BSSs when the length of each fragment is between lengths 50 and 10 base pairs, with some BSSs dropping heavily in performance between 100 and 50 base pairs for specific datasets (e.g. Sequence 8.1 for SeqScreen). Although in both SeqScreen and Commec, lower lengths below 50 base pairs that are flagged have a high percentage that do not classify as the specific toxin or agent’s taxonomic ID, but is still flagged. In this study, we still consider these fragments as successfully flagged due to the BSSs recognizing that the fragment contains a SOC.

These tables also show how skewed the averages are as the point where performance drops are typically past half of the lengths we present (300 to 10 base pairs).

Table 4 shows that when encapsulated, some toxins and agents cannot be classified by any BSSs we evaluate. Specifically, all BSSs have trouble identifying Sequence 1; this may be due to the small size of our Sequence 1 sample which is only the chain A of the larger genome (see Table 1).

**TABLE 4.**
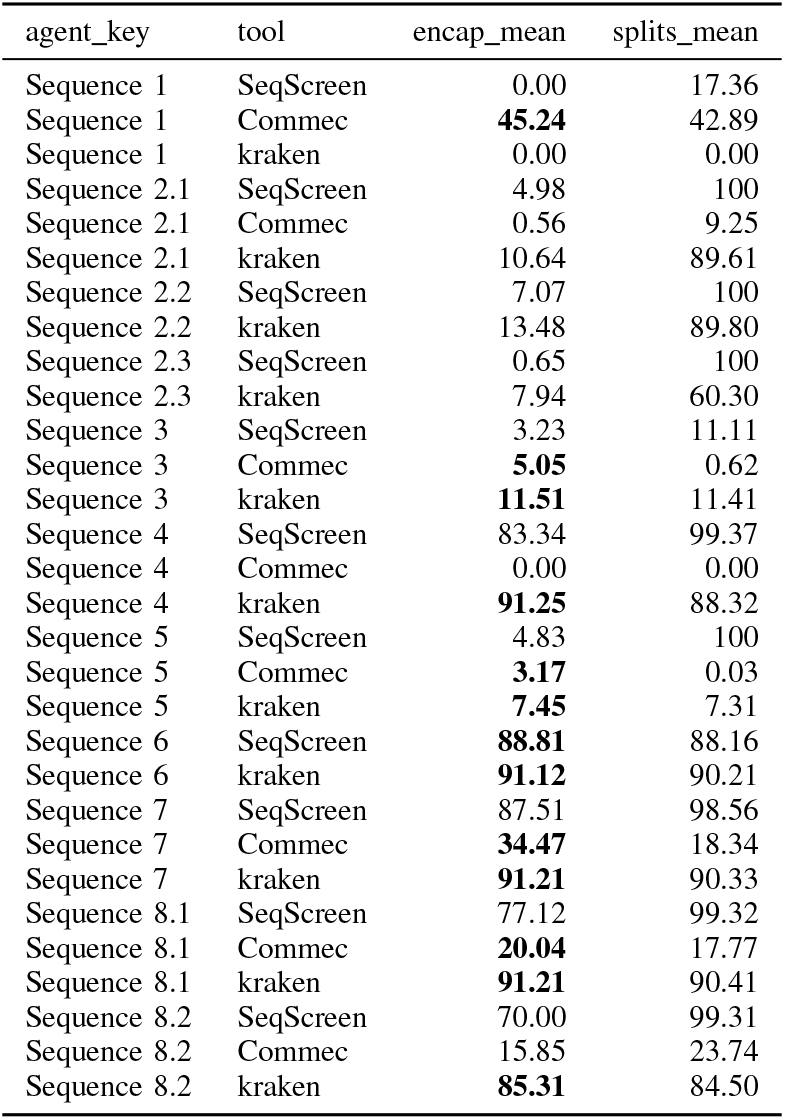
Average detection rate for encapsulation and splits .

**TABLE 5.** Detection rates at every 50 lengths for Seq Screen evaluating encapsulated data.

| dataset | L300 | L250 | L200 | L150 | L100 | L50 | L10 |
| --- | --- | --- | --- | --- | --- | --- | --- |
| Sequence 1 | 0.00 | 0.00 | 0.00 | 0.00 | 0.00 | 0.00 | 0.00 |
| Sequence 2.1 | 6.11 | 6.226 | 6.17 | 6.10 | 5.93 | 3.78 | 0.83 |
| Sequence 2.2 | 8.60 | 9.04 | 8.91 | 9.04 | 8.76 | 4.59 | 0.79 |
| Sequence 2.3 | 1.15 | 1.11 | 1.05 | 0.92 | 0.28 | 0.024 | 0.01 |
| Sequence 3 | 0.00 | 1.20 | 1.32 | 1.27 | 5.386 | 8.790 | 1.61 |
| Sequence 4 | 100 | 100 | 100 | 100 | 99.47 | 88.86 | 24.03 |
| Sequence 5 | 0.00 | 6.48 | 6.66 | 6.84 | 6.73 | 0.0000 | 0.01 |
| Sequence 6 | 100 | 100 | 100 | 100 | 99.48 | 91.09 | 21.04 |
| Sequence 8.1 | 100 | 100 | 100 | 100 | 98.85 | 42.95 | 5.12 |
| Sequence 8.2 | 100 | 98.98 | 96.16 | 88.98 | 81.02 | 0.00 | 1.95 |

**Table 6.** Detection rates at every 50 lengths for Commec evaluating encapsulated data.

| dataset | L300 | L250 | L200 | L150 | L100 | L50 | L10 |
| --- | --- | --- | --- | --- | --- | --- | --- |
| Sequence 1 | 100 | 100 | 100 | 57.80 | 2.400 | 0.00 | 0.00 |
| Sequence 2.1 | 2.15 | 1.57 | 0.90 | 0.47 | 0.22 | 0.013 | 0.022 |
| Sequence 3 | 0.00 | 11.79 | 8.98 | 6.20 | 2.19 | 0.00 | 0.00 |
| Sequence 4 | 0.00 | 0.00 | 0.00 | 0.00 | 0.00 | 0.00 | 0.00 |
| Sequence 5 | 0.00 | 0.00 | 0.00 | 8.25 | 7.44 | 1.38 | 0.10 |
| Sequence 8.1 | 38.10 | 37.38 | 34.18 | 26.75 | 13.04 | 3.63 | 0.68 |

As a preliminary specificity check, we evaluated a Aequorea victoria green-fluorescent (GFP) gene as a benign control sequence and observed no false positives across both Commec and SeqScreen. However, we acknowledge that this represents a limited assessment of specificity. In particular, evaluating benign sequences under transformations similar to those used in our adversarial setting is an important direction for future work to better characterize false positive behavior in realistic screening scenarios.

We note that failure to flag a sequence should not be interpreted as evidence of safety, as biological risk can depend on downstream assembly, expression context, host system, and combination with other sequence elements.

### 6.1. Direct BSS Comparisons

In Figure 4, we can pinpoint which specific fragments SeqScreen is failing to identify. We can generate these analytics for any BSS. The y-axis is the length of the fragment and the x-axis is the position of the genome of Sequence 8.1. Each visible block is a fragment of a specific length determined by the y-axis that was run on SeqScreen. The further up the graph is, the larger the “blocks” will be. Notably we can see a clear drop in performance when the size of the fragments drops below a certain point similar to Figure 3. For Sequence 8.1, SeqScreen struggles below a size of 80 base pairs. We can further look into Sequence 8.1 in Figure 4. We notice that at times SeqScreen can be inconsistent, where a fragment can be marked as malicious but later a similar region of the genome can be marked as benign for a smaller fragment. This analysis can also be used to pinpoint vulnerable areas in a sequence. The annotations on the figure display the specific genes and the location of the gene with respect to the entire genome. With the annotations we can see that SeqScreen is specifically strong on proteins like A, G, and E. SeqScreen is able to detect these genes as malicious past the typical point where SeqScreen drops in performance. Because of the difference, we can clearly see how SeqScreen treats each gene. Similar to the BSSs, Kraken also fails to classify once the fragment size is too small. But we cannot view this comparison directly as Kraken has a different goal than SeqScreen does, hence we compare SeqScreen to Commec.

Figure 5 shows most datasets that are simply split achieve a high detection rate with SeqScreen at most lengths. However, we see a unique pattern where specifically Sequence 2.2, Sequence 4, Sequence 6, Sequence 7, and Sequence 8.1, all drop off in performance at a similar length range. Sequence 3 is also unique where the performance of SeqScreen increases as lengths become smaller.

Figure 6 shows a direct comparison between SeqScreen and Commec over Sequence 8.1 and Sequence 2.1 positionally. In this analysis, each data point is a fragment of the respective genome. Thus, the lower the length the smaller the fragment. In Sequence 8.1, SeqScreen scores 32.7% better than Commec with SeqScreen achieving 40.2% detection rate while Commec achieves a 7.5% detection rate. With fragments with lengths above 50 base pairs, SeqScreen widens the gap by outperforming Commec by 62.4% with SeqScreen achieving a 77.2% detection rate while Commec only achieving a 14.8% detection rate.

#### 6.1.1. Annotations

We also provide annotations for Sequence 8.1 in Figure 4 and Figure 6 that show an annotation for each gene labeled and the resulting protein labeled as letters so as not to reveal what Sequence 8.1 is. The Figures show SeqScreen is particularly strong at detecting protein A (a polymerase complex protein), protein B (membrane-associated protein), and protein C (spike glycoprotein). SeqScreen detects these proteins at shorter fragment lengths than other portions of the genome. In Sequence 8.1, Commec detects well until fragment lengths drop below 100 bp, whereas SeqScreen can detect these genes in even shorter lengths. SeqScreen has minor weakness detecting parts of L (RNA-dependent RNA polymerase). We can see small gaps in covers, covers-a, and covers-random indicating these specific regions show weakness as manipulation complexity increases. Our insights do not show whether a fragment should be flagged as malicious, simply that the BSS flagged the fragment as malicious.

We can also see that SeqScreen has weaknesses in the gaps between each gene when we get to the covers-a and covers-random dataset, often not detecting the gaps between the genes at all even when the fragment lengths are at 300 base pairs. However, Commec excels at finding the gaps between each of the genes and fails at the genes themselves, in an opposite manner of SeqScreen. Of course, Commec in general does not perform well. Commec does flag a lot of fragments in B, D, and E, but not to the rate of SeqScreen. Commec has trouble identifying the NP (Nucleoprotein), C (Spike Glycoprotein), and L (RNA-dependent RNA polymerase or “Large” Protein). The RNA-dependent RNA polymerase does not work without A and the nucleoprotein. Not being able to identify the spike protein of Sequence 8.1 raises a significant concern. The spike protein in any viral compound is responsible for the viral agent to attach to host cells of the infected and mediating membrane fusion to allow for viral entry. If an adversary were aware that a BSS is specifically weak at detecting spike proteins, they can isolate this specific spike protein and attempt to get this spike protein to work on a different viral agent that could be more difficult for a screener to detect.

SeqScreen does maintain a high detection rate up to fragments with a length of 50 when SeqScreen does recognize the agent. Table 5 shows when SeqScreen does identify the toxin or agent, the detection rate is consistent as lengths get shorter, for example Sequence 4 maintains 100% detection rate up to when the split lengths are 100, where SeqScreen’s detection rate drops by less than 1%. Then the detection rate falls drastically between lengths 50 and 10. We can see in Figure 3 the performance drop happens around lengths 25 to 30.

### 6.2. Kraken

Although Kraken is not a BSS, we felt Kraken would provide a good baseline for BSSs. Kraken is a bioinformatics tool for rapidly assigning taxonomic labels to DNA sequences. The objective of the tool is to label sequences and not to determine whether a sequence is malicious. Therefore, all results are based on whether Kraken correctly classified the taxonomic label for the sequences. If a taxonomic label was close to the sequence, we accepted that as a true positive result for Kraken. We determine if a label is close by manually selecting related taxonomic ids in NCBI. We choose to evaluate Kraken because the field of taxonomic identification is more mature than BSSs. Kraken also has an easier task than BSSs do, giving us an idea of what we want a BSS in the future to look like.

Figure 7 shows Kraken maintains performance until Kraken attempts to evaluate fragments of lengths less than 25. And compared to Figure 3, the drop off in performance is steeper but happens in similar places despite the tools having different purposes. A large difference between the Kraken and the BSSs is the point where performance reaches a near 0% detection rate when similar to the BSSs. There are some agents that Kraken does not recognize and most likely is missing from the standard database. Kraken does have guides on how to create and add to the existing database.

**Figure 7.**
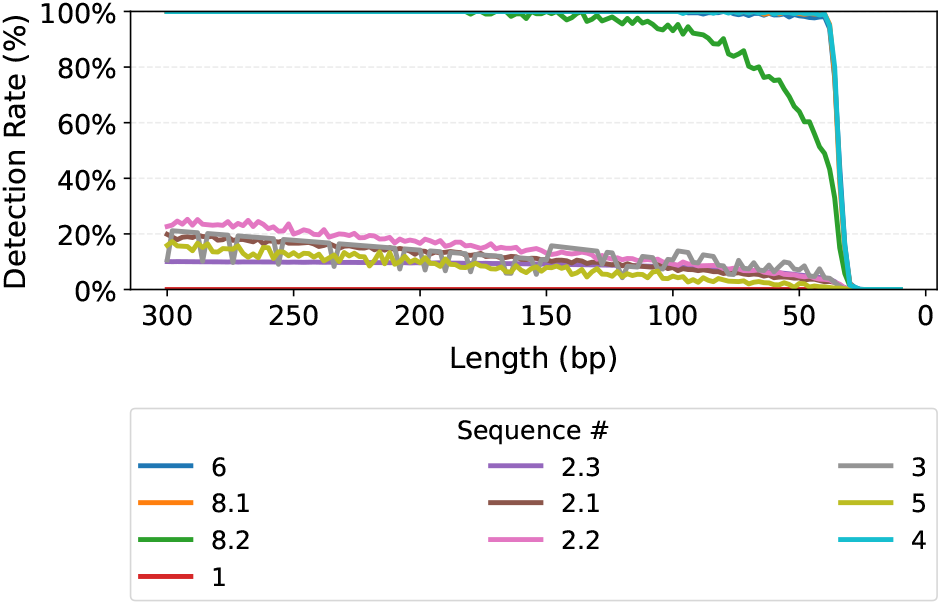
Detection rate at each split length for each agent in Kraken.

### 6.3. Time Analysis

In addition to detection rate, we compare the runtime of each BSS. Table 7 demonstrates the differences among the BSSs. Commec takes the longest by far averaging 11 hours per complete analysis of Sequence 5, followed by SeqScreen which averages approximately a few hours, and finally Kraken which takes only a few milliseconds. The Kraken results are expected due to Kraken not being a BSS but rather a classification model, meaning Kraken does not have to determine whether a sequence is malicious, skipping an entire classification step. However, the differences between SeqScreen and Commec are significant with respect to their performance. Commec takes significantly longer than SeqScreen due to Commec aligning the input sequence using BLAST [33] and DIAMOND [32] while SeqScreen instead attempts to classify the sequence using a deterministic model and seeking out Sequence-of-Concerns (SOCs) using a deterministic model. Table 7 also shows the average times in more detail along with the size of each dataset of the respective toxin or agent. The time taken for each dataset is almost proportional to the size of the dataset only in Commec while in SeqScreen and Kraken, Sequence 2.1 has a longer average runtime than Sequence 5 despite being shorter. We suspect this is because of Commec’s pipeline of running BLAST multiple times regardless before determining if a sequence may be malicious.

**TABLE 7.** Average run -time (s) of different BSSs across different datasets relative to size (kb).

| BSS | dataset | mean(s) | size(kb) |
| --- | --- | --- | --- |
| Commec | Sequence 5 | 39650 | 2100 |
| Commec | Sequence 3 | 11500 | 282.5 |
| Commec | Sequence 2.1 | 7159 | 181.9 |
| Commec | Sequence 6 | 3223 | 19.10 |
| Commec | Sequence 8.1 | 3265 | 19.00 |
| Commec | Sequence 7 | 2743 | 11.70 |
| SeqScreen | Sequence 5 | 570.6 | 2100 |
| SeqScreen | Sequence 3 | 307.5 | 282.5 |
| SeqScreen | Sequence 2.1 | 505.8 | 181.9 |
| SeqScreen | Sequence 6 | 253.7 | 19.10 |
| SeqScreen | Sequence 8.1 | 251.4 | 19.00 |
| SeqScreen | Sequence 7 | 233.8 | 11.70 |
| Kraken | Sequence 5 | 9.280 | 2100 |
| Kraken | Sequence 3 | 13.34 | 282.5 |
| Kraken | Sequence 2.1 | 13.86 | 181.9 |
| Kraken | Sequence 8.1 | 8.670 | 19.00 |
| Kraken | Sequence 6 | 9.670 | 19.10 |
| Kraken | Sequence 7 | 9.670 | 11.70 |

## 7. Discussion

Here we discuss how our benchmark can find weaknesses in BSSs, so developers can pinpoint where to focus problem-solving efforts and to interpret the findings of our experiments outlined in Section 6.

Overall performance as defined in Table 3 shows most open-source BSSs cannot classify manipulated input at a rate above a 50% detection rate. Commec mentions that the first major focus is to classify plain adversarial data first and an adjacent work [6] mentions that manipulation testing is still being developed and has yet to be explored in depth. Commec launched the first official version in August of 2025, but launched the first public alpha version in May 2024. Meanwhile SeqScreen was launched in 2020 and is also still in active development.

Table 3 shows that manipulation of fragments can greatly affect BSS performance. We can also see this in Figure 3 and Figure 5 where after each fragment is encapsulated, SeqScreen performance drops much earlier at larger lengths than with non-manipulated splits.

We find that attackers can pinpoint specific split lengths to bypass BSSs regardless of manipulation. In Figure 5, SeqScreen performance on a group of datasets drops heavily between the 20 to 30 base pairs and comes back up in performance, offering a narrow window for adversaries to use. With encapsulation in Figure 3 and other manipulations, adversaries can simply split sequences small enough to bypass SeqScreen.

With manipulations, BSS performance generally drops (Table 3) by an average of 28% between splits and encapsulation. Demonstrating manipulations can become a large problem for BSSs. However, not all manipulations cause a decrease in performance. In Table 4, the bolded rows are where encapsulation results in an increase of average detection rate. In cases where encapsulation improves detection, the average increase is only about 2.73%. A minuscule difference compared to when manipulations are introduced. However, this demonstrates Commec’s and Kraken’s resilience to manipulations and suggests possibilities for formatting fragments to increase detection rates in BSSs.

We would like to emphasize if a BSS does not flag a high percentage of the data, this may not necessarily indicate the BSS is performing poorly. In Figure 4, we can see gene L, the RNA-dependent RNA polymerase, is flagged up to around 80 base pairs. However, in Figure 6, we can see Commec does not flag most of L. Whether the RNAdependent RNA polymerase should be flagged is debatable. The gene is shared among other genomes that may not be malicious, however, in this case the gene is associated with the malicious agent. Our work and framework is not intended to determine whether or not a fragment or gene should be flagged, but rather to encourage discussion on whether it should be flagged. Different parties trying to implement BSSs into their pipeline should determine whether a specific BSSs fits the pipeline logistically, our framework helps with that processes by comparing and looking into BSS behaviors.

Despite Commec taking more time and resources, SeqScreen proves to be the more accurate BSS when flagging manipulated sequences. Commec has a nearly 1TB database compared to SeqScreen’s 180GB database. Commec uses BLAST and DIAMOND to do extensive analysis and alignment that sometimes takes more than an hour to perform a single run while SeqScreen can do a run in a matter of minutes. In Table 5 we can see SeqScreen performed well with Sequence 4, the Sequence 6, and both forms of Sequence 8. And we can see those detection rates are maintained within a reasonable window until the lengths are below 50. But SeqScreen does not classify Sequence 1, Sequence 3, and Sequence 5 at all. A plausible explanation is that SeqScreen is better suited to manipulated short fragments because it was designed for functional characterization of short sequences via curated FunSoCs and complementary evidence sources, whereas Commec is primarily a synthesis-screening pipeline that relies on profile- and homology-based screening. Both BSSs target synthetic-sequence screening, but they were not principally presented as defenses against adaptive adversaries. SeqScreen was explicitly designed for short DNA sequences and oligonucleotides, while Commec is positioned as a provider-oriented synthesis-order screening baseline.

Our metrics in this study heavily penalize misclassifications of a single fragment relative to how other studies view BSSs. In a scenario where an attacker attempts to submit all the fragments of a genome at a specific length at once, only one fragment needs to be flagged for the whole order to be suspicious. Or the attacker could split up each fragment order over a period of time to try to hide intention. To combat this, a system could be developed to monitor order history every time a new order is placed. However, an attacker could circumvent a system by creating new accounts for each fragment or sending each fragment to a different provider as described in Figure 1, making independent fragment detection crucial to BSSs. An attacker could also select fragments they know can bypass screeners. This is less realistic because the missing regions would still need to be obtained and integrated to produce a functional construct. But this is still a scenario we should prepare for.

We use Kraken in this study as a comparison to BSSs because Kraken has an easier task and is a more mature tool. Table 7 demonstrates that Kraken also cannot identify manipulated Sequence 1 fragments and performs poorly on Sequence 5. Table 3 shows that Kraken beat Commec in on average but only beats SeqScreen in encapsulation, and then drops in average performance as manipulations become more complex. Showing that SeqScreen is more resilient to manipulated datasets than both Commec and Kraken. Bringing questions and further review to how both BSS and taxonomic classifiers draw information from a database.

Overall, we find there needs to be more security evaluations on BSSs. In this study we only tested open-source BSSs and we are unaware if private BSSs perform better or worse. Each BSS has different methods to identify sequences resulting in varied results for each BSS. However, because they are not open source, we cannot evaluate them. Due to synthetic nucleic acid and protein technology being available to the public now, we need to push for collaborative testing to be done by the community rather than internal testing teams.

## 8. Conclusion

We believe that additional work and testing are needed to better secure synthetic nucleic acid screening systems and benchmarking systems, such as the benchmark we design and explore here, can have a valuable role in such work.

Under our benchmark, across each length category treated as a continuous ordering of fragments, SeqScreen achieved a 79.27% detection rate, while Commec achieved an 18.67% detection rate. Current open-source BSSs may perform adequately against unsophisticated (“lazy”) adversaries, and possibly against automated adversaries. However, to prepare for real bioterrorism threats, we must evaluate these systems against motivated and capable adversaries.

As shown above, the BSSs we tested achieved only 43% and 11% success rates in detecting dangerous toxins and agents such as Sequence 8.1 and Sequence 2.1. These percentages are influenced by toxins and agents that the BSSs cannot detect at all. When SeqScreen is able to detect a sequence, its performance remains relatively robust until fragments become sufficiently short that general genomic classifiers can no longer reliably classify them.

Our data suggest that if an adversary can recombine fragments in a laboratory setting, they could assemble a wide range of select toxins and agents with little to no chance of detection. Although some BSSs explicitly acknowledge that detection efficiency decreases below certain length thresholds, more systematic testing is needed to precisely characterize these thresholds. In the meantime, manufacturers should consider complementary safeguards beyond BSS-based screening until these limitations are addressed.

Stepping back, we believe it is valuable to benchmark potentially dangerous capabilities across increasing levels of complexity, reflecting both simple and sophisticated realworld attack strategies. Evaluations that directly measure how BSS limitations could enable harmful outcomes can serve as early warning signals and guide the development of more robust biosecurity safeguards.

## Acknowledgments

We thank the anonymous reviewers and our shepherd, David Molik, for their invaluable feedback and guidance during the revision process. We are also grateful to the University of Washington Security and Privacy Research Lab (including Grace Brigham, Ian Chang, Kshitish Ghate, Gregor Haas, Alexandra Michael, Rachel McAmis, and Franzi Roesner) for insightful feedback. This project was supported in part by the University of Washington Tech Policy Lab and by the University of Washington Molecular Information Systems Lab. Henry C. Wong was additionally supported in part by the University of Washington College of Engineering Dean’s Fellowship. Tadayoshi Kohno is supported by the Robert L. McDevitt, K.S.G., K.C.H.S. and Catherine H. McDevitt L.C.H.S. Chair in Computer Science at Georgetown University.

## Appendix A

### Ethical Considerations

Affected parties in our work include researchers and developers of open-source and closed-source Biosecurity Screening Software (BSS). Our study focuses only on open-source software while leaving closed-source for future studies. Companies that manufacture synthetic DNA and consumers that order synthetic DNA (e.g. research laboratories and drug manufacturers) are also direct stakeholders as BSSs add an extra step in processing orders for synthetic DNA and affects a consumer’s accessibility to synthetic DNA services. Our work also affects governments and policy makers as there have been recent pushes on securing nations against bio-attacks and synthetic DNA manufacturing regulation. Lastly, our own research teams are stakeholders as our own research labs frequently order synthetic DNA for our own experiments.

We acknowledge there may be potential misuse of our data. Malicious actors could use our methods to actively bypass BSSs to perform bio-attacks.

To mitigate negative impacts we share our data with the developers of the BSSs that we evaluated in our work and will provide our data to any BSS developer that requests the data. We first provide BSS developers with our input data, malicious sequences that have been manipulated as if an adversary were attempting to bypass BSSs. This data stress-tests and benchmarks the BSS against potential adversarial orders. We also provide software to generate these manipulations if the BSS developer wishes to test toxins and agents we did not review. We then offer a range of analytics scripts that allows developers to view specifics on where the weak points of the BSS are. Our work also offers one of the first looks into how open-source BSSs perform against manipulated and split genomes. We do not disclose which agents and toxins we used to benchmark the BSSs. We do not open-source our manipulated data and make it available only by request.

The objective of this work is to present a benchmark for BSSs and to promote further work in securing BSSs. We have notified the developers of Commec and SeqScreen and have been in discussion with both developers on how to better improve BSSs. SeqScreen has notified us of a new version that should improve BSS performance. We considered the risk associated with adversaries learning how to attack BSSs but came to the conclusion that executing an attack from our public data is not trivial. As in prior work [25], we use the same practice and mention the names of the software. DNA synthesis manufacturers also typically do not disclose which BSS they use. Thus, releasing the names does not assist adversaries in their attack on individual synthesis companies. We did not identify any ethical concerns sufficient to warrant withholding publication given the mitigation and usage described above. We believe open-sourcing our work will promote further exploration of how adversaries can bypass BSSs and help build better BSSs.

## Appendix B

### Manipulation Implementation

#### B.1. Splits

Given an input nucleotide sequence *S*, we generate fragmented datasets by choosing a target fragment length *k* and partitioning *S* into contiguous, non-overlapping sub-sequences of length at most *k*. For each *k*, fragments are extracted as *S*[*ik* : min((*i*+1)*k* − 1, |*S*|)] for *i* = 0, 1, 2, …. If incomplete fragments are excluded, any terminal fragment shorter than *k* is discarded; otherwise, it is retained. We repeat this procedure over a range of fragment lengths, producing a separate output set for each *k*.

Below is a simplified pseudocode version of our script to split each sequence.

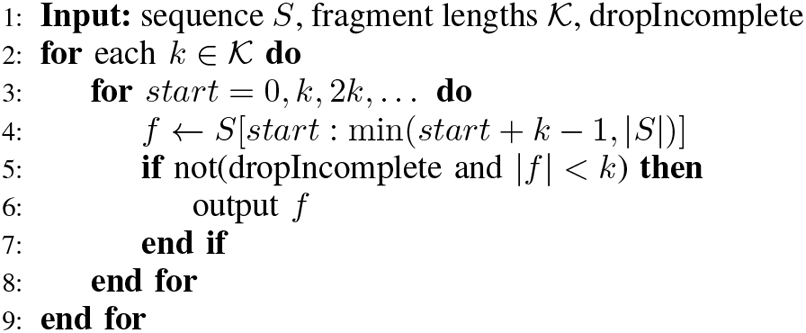

#### B.2. Encapsulation

Given a fragment *f* of length *k*, we generate an encapsulated fragment by adding random flanking sequences to both ends of the fragment. Specifically, we sample a left flank *l* and a right flank *r*, each of fixed length 4 bp, uniformly at random from {*A, C, G, T*}4, and construct the modified sequence as *f ′* = *l* ∥*f*∥ *r*. We apply this transformation independently to every fragment in the fragmented dataset. This manipulation serves as a simplified proxy for common ordering formats in which the core target sequence is submitted with short additional context, such as adapters, primers, or assembly-related ends.

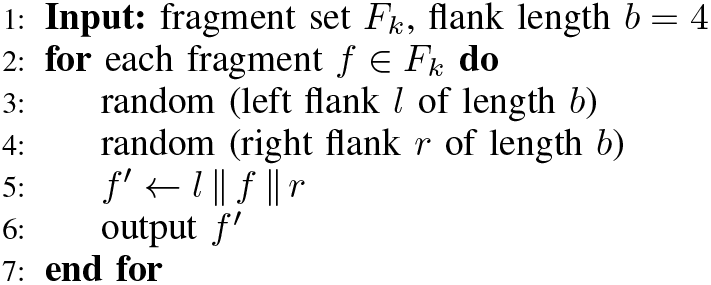

#### B.3. Flipped Splits

Given an input nucleotide sequence *S* of length *L*, we generate a flipped sequence by introducing random point mutations at a subset of positions. For a specified mutation budget *n* (or equivalently a mutation percentage), we sample *n* distinct positions uniformly without replacement from {1, …, *L*}. At each selected position, the original nucleotide is replaced with a uniformly chosen alternative base from {*A, C, G, T*} \ {*Si*}, ensuring that every selected position is changed. This manipulation is intended as an algorithmic stressor for sequence-matching approaches rather than as a biologically realistic model of low-effort adversarial construction.

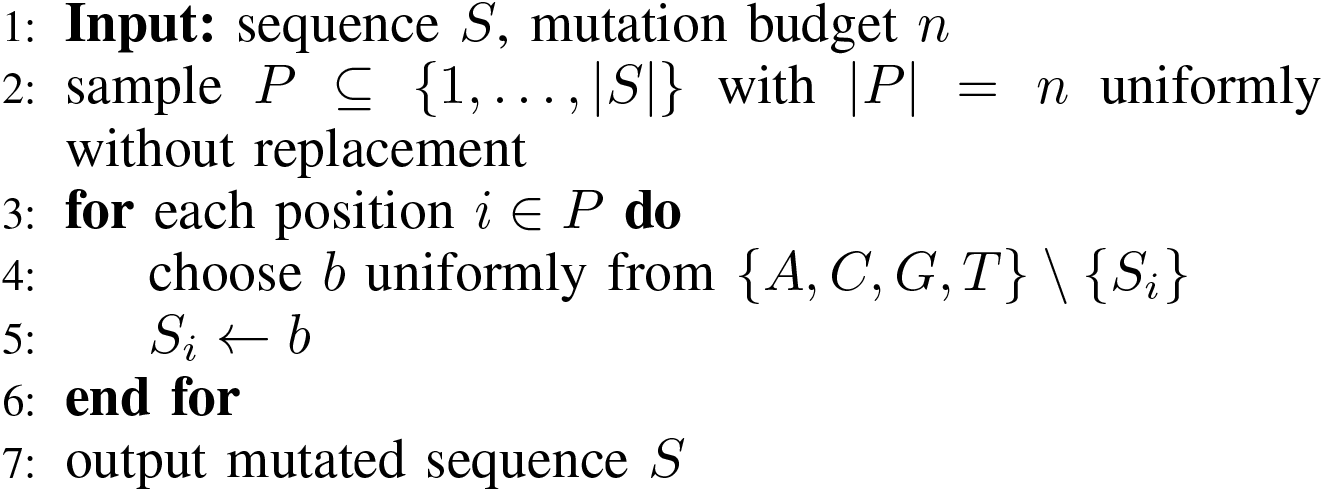

#### B.4. Covers

Given an input fragment *f* of length *k*, we generate a covered fragment by appending auxiliary sequence context derived from a benign donor sequence. Specifically, we set the appended length to *c* = ⌈1.5*k*⌉ and construct the modified sequence as *f*′ = *f* ∥ *d*[1 : *c*], where *d* is the donor sequence. When the donor sequence is shorter than *c*, we cycle the donor sequence as needed until the required appended length is reached. In our experiments, the donor sequence for *covers* is a GFP gene. This manipulation is intended to test whether appending sufficiently large benign context can shift screener behavior away from the malicious fragment.

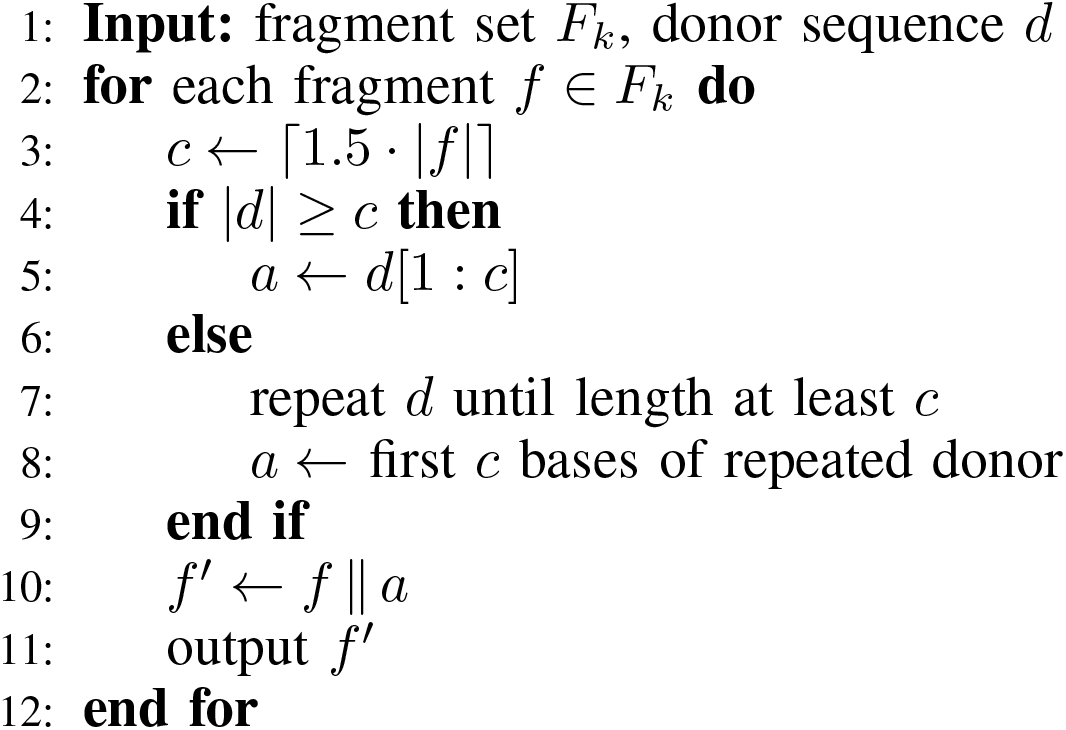

## References

[1] N. S. C. on Emerging Biotechnology (NSCEB), “Charting the future of biotechnology: An action plan for american security and prosperity.” https://www.biotech.senate.gov/final-report/chapters/, Apr. 2025. Accessed: 2026-02-02.

[2] Office of Science and Technology Policy and Administration for Strategic Preparedness and Response, “Framework for nucleic acid synthesis screening,” tech. rep., U.S. Government, September 2024. Accessed: 2025-07-30.

[3] Senate Select Committee on the Future of Biotechnology, “Section 4: Oversight and security,” 2024. Part of the Final Report of the Senate Bipartisan Committee on Biotechnology. Accessed: 2025-07-30.

[4] Department of Health and Human Services, “Screening framework guidance for providers of synthetic double-stranded dna.” Federal Register, Oct. 2010. Document No. 2010-25728.

[5] U.S. Department of Health and Human Services, “Screening framework guidance for providers and users of synthetic nucleic acids,” tech. rep., Administration for Strategic Preparedness and Response (ASPR), U.S. Department of Health and Human Services, 2023. Accessed: 2026-02-28.

[6] N. E. Wheeler, C. Bartling, S. R. Carter, A. Clore, J. Diggans, K. Flyangolts, B. T. Gemler, B. Rife Magalis, and J. Beal, “Progress and prospects for a nucleic acid screening test set,” Applied Biosafety, vol. 29, no. 3, pp. 133–141, 2024.

[7] B. T. Gemler, C. Mukherjee, C. Howland, P. A. Fullerton, R. R. Spurbeck, L. A. Catlin, A. Smith, A. T. Minard-Smith, and C. Bartling, “Ultraseq, a universal bioinformatic platform for information-based clinical metagenomics and beyond,” Microbiology Spectrum, vol. 11, no. 3, pp. e04160–22, 2023.

[8] J. Beal, D. Wyschogrod, T. Mitchell, S. Katz, J. Manthey, and A. Clore, “Development and transition of fast-na screening technology,” Tech. Rep. BBN Tech Report 8622, 2021.

[9] N. E. Wheeler, S. R. Carter, T. Alexanian, C. Isaac, J. Yassif, and P. Millet, “Developing a common global baseline for nucleic acid synthesis screening,” Applied Biosafety, vol. 29, no. 2, pp. 71–78, 2024.

[10] A. Balaji, S. Dey, D. Vyshedskiy, B. Hall, C. Shih, C. Liu, W. O. Osburn, J. Diggans, A. Gitter, and M. Pop, “Seqscreen: accurate and sensitive functional screening of pathogenic sequences via ensemble learning,” Nature Communications, vol. 13, no. 1, p. 5204, 2022.

[11] K. E. Rey Edison, Shay Toner, “Evaluating the robustness of current nucleic acid synthesis screening.” https://drive.google.com/file/d/1hNUnU8i2yubt5uesmmV17aTJXhYYDgTY/edit, 2024. Unpublished manuscript. Available at: https://drive.google.com/file/d/1hNUnU8i2yubt5uesmmV17aTJXhYYDgTY/edit.

[12] B. J. Wittmann, T. Alexanian, C. Bartling, J. Beal, A. Clore, J. Diggans, K. Flyangolts, B. T. Gemler, T. Mitchell, S. T. Murphy, et al., “Strengthening nucleic acid biosecurity screening against generative protein design tools,” Science, vol. 390, no. 6768, pp. 82–87, 2025.

[13] International Gene Synthesis Consortium (IGSC), “Harmonized screening protocol v3.0,” tech. rep., International Gene Synthesis Consortium, september 2024. Accessed: 2026-02-28.

[14] J. Diggans and E. Leproust, “Next steps for access to safe, secure dna synthesis,” Frontiers in bioengineering and biotechnology, vol. 7, p. 86, 2019.

[15] S. Kosuri and G. M. Church, “Large-scale de novo DNA synthesis: technologies and applications,” Nature Methods, vol. 11, no. 5, pp. 499–507, 2014.

[16] R. A. Hughes and A. D. Ellington, “Synthetic DNA synthesis and assembly: putting the synthetic in synthetic biology,” Cold Spring Harbor Perspectives in Biology, vol. 9, no. 1, p. a023812, 2017.

[17] M. G. Ryadnov, “DNA synthesis technologies to close the gene writing gap,” Nature Reviews Chemistry, vol. 7, pp. 144–161, 2023.

[18] D. G. Gibson, L. Young, R.-Y. Chuang, J. C. Venter, C. A. Hutchison III, and H. O. Smith, “Enzymatic assembly of DNA molecules up to several hundred kilobases,” Nature Methods, vol. 6, no. 5, pp. 343–345, 2009.

[19] D. DiEuliis, S. R. Carter, and G. K. Gronvall, “Options for synthetic DNA order screening, revisited,” mSphere, vol. 2, no. 4, pp. e00319–17, 2017.

[20] National Institute of Standards and Technology, “NIST test dataset for assessing baseline nucleic acid sequence screening,” 2025.

[21] T. S. Laird, K. Flyangolts, C. Bartling, B. T. Gemler, J. Beal, T. Mitchell, S. T. Murphy, J. Berlips, L. Foner, R. Doughty, et al., “Inter-tool analysis of a nist dataset for assessing baseline nucleic acid sequence screening,” Applied Biosafety, p. 15356760251401228, 2025.

[22] N. E. Wheeler, “Responsible ai in biotechnology: balancing discovery, innovation and biosecurity risks,” Frontiers in Bioengineering and Biotechnology, vol. 13, p. 1537471, 2025.

[23] Y. Chen, M. Tucker, N. Panickssery, T. Wang, F. Mosconi, A. Gopal, C. Denison, L. Petrini, J. Leike, E. Perez, and M. Sharma, “Enhancing model safety through pretraining data filtering,” Aug. 2025.

[24] OpenAI, “Openai’s approach to frontier risk,” Oct. 2023.

[25] S. Tayouri, V. Kogan, J. Beal, T. Levy, D. Farbiash, K. Flyangolts, S. T. Murphy, T. Mitchell, B. Rotblat, I. Vaksler-Lublinsky, et al., “Defending synthetic dna orders against splitting-based obfuscation,” bioRxiv, pp. 2025–03, 2025.

[26] A. S. Rathore, S. Choudhury, A. Arora, P. Tijare, and G. P. Raghava, “Toxinpred 3.0: An improved method for predicting the toxicity of peptides,” Computers in biology and medicine, vol. 179, p. 108926, 2024.

[27] C. Baum, J. Berlips, W. Chen, H. Cozzarini, H. Cui, I. Damgård, J. Dong, K. M. Esvelt, L. Foner, M. Gao, et al., “A system capable of verifiably and privately screening global dna synthesis,” National Science Review, p. nwag103, 2026.

[28] D. C. Ince, L. Hatton, and J. Graham-Cumming, “The case for open computer programs,” Nature, vol. 482, no. 7386, pp. 485–488, 2012.

[29] Federal Select Agent Program, “HHS and USDA select agents and toxins,” tech. rep., U.S. Department of Health and Human Services and U.S. Department of Agriculture, january 2025. Accessed: 2026-02-28.

[30] S. Federhen, “The ncbi taxonomy database,” Nucleic acids research, vol. 40, no. D1, pp. D136–D143, 2012.

[31] D. E. Wood, J. Lu, and B. Langmead, “Improved metagenomic analysis with kraken 2,” Genome biology, vol. 20, no. 1, p. 257, 2019.

[32] B. Buchfink, K. Reuter, and H.-G. Drost, “Sensitive protein alignments at tree-of-life scale using diamond,” Nature methods, vol. 18, no. 4, pp. 366–368, 2021.

[33] C. Camacho, G. Coulouris, V. Avagyan, N. Ma, J. Papadopoulos, K. Bealer, and T. L. Madden, “Blast+: architecture and applications,” BMC bioinformatics, vol. 10, no. 1, p. 421, 2009.

